# Myo1e/f at the podosome base regulate podosome dynamics and promote macrophage migration

**DOI:** 10.1101/2025.04.28.651090

**Authors:** Pasquale Cervero, Sarah R. Barger, Perrine Verdys, Robert Herzog, Tyler Paul, Marjorie Palmieri, Renaud Poincloux, Stefan Linder, Mira Krendel

## Abstract

Cells of the monocyte lineage form specialized membrane-associated, actin-rich structures, called podosomes. Podosomes play important roles in cell adhesion and migration as well as the proteolytic degradation of the extracellular matrix. While podosomes are always closely associated with the plasma membrane, the structural components linking the podosome core, composed of branched actin, to the membrane are not fully understood. In this study we show that class I myosins, Myo1e and Myo1f, localize to a specific region of podosomes, underneath the podosome core and near the ventral plasma membrane, and that this localization is mainly mediated by the Myo1e/f TH2 domains. Respective knockdowns or knockouts of Myo1e/f lead to increased podosome size, altered turnover and lateral mobility, which is likely due to Myo1e/f regulating the attachment of core actin filaments to the plasma membrane. In addition, Myo1e/f double knockout macrophages were characterized by a reduction in 3D and 2D migration, even though these cells exhibited increased ability to degrade the extracellular matrix. Along with the other membrane-associated podosome components, such as the transmembrane proteins MT-MMP and CD44, and the GPI-anchored DNase X, Myo1e and Myo1f mark the membrane-proximal region of podosomes. We propose to label this region as the podosome “base”, an additional substructure joining the current trifecta of the podosome cap, core, and ring.

## Introduction

Macrophages are cells of the innate immune system that actively migrate along the surfaces of endothelial cells, across the endothelial and basement membrane barriers, and throughout tissues and interstitial spaces to perform their immune surveillance functions. They are highly dynamic cells that can employ a variety of cell migration modalities. This migratory behavior is often reliant on the formation of podosomes, specialized actin-rich adhesion structures, on their ventral surfaces (Wiesner et al., 2014). In addition to forming cell-matrix adhesion sites, podosomes also serve as the sites of proteolytic enzyme secretion, allowing macrophages to invade and breach barriers (Linder et al., 2023).

Podosomes consist of >300 different proteins (Cervero et al., 2012) that form a characteristic tripartite organization involving an F-actin-rich core surrounded by a ring of adhesion proteins and topped by a cap structure (Linder et al., 2023). The F-actin core of podosomes includes the Arp2/3 complex (Linder et al., 2000) that promotes assembly of branched actin networks, Arp2/3 activators belonging to the WASp protein family (Linder et al., 1999), actin cross-linker L-plastin (Zhou et al., 2016), as well as the actin-binding protein cortactin (Tehrani et al., 2006) and Tks family scaffold proteins (Seals et al., 2005), among others. The discontinuous ring structure contains proteins such as integrins (Marchisio et al., 1988), paxillin (DeFife et al., 1999), and vinculin (DeFife et al., 1999; Marchisio et al., 1988), while the cap structure contains mostly actin crosslinking or bundling proteins such as α-actinin (van den Dries et al., 2019), fascin (Van Audenhove et al., 2015), or LSP1 (Cervero et al., 2018). Altogether these proteins cooperate to regulate podosome life cycle and function, but the role of many podosome components remains to be elucidated.

Myosin 1e (Myo1e) and the closely related myosin I isoform myosin 1f (Myo1f) were first identified as podosome components by mass spectroscopy of podosome-enriched fractions from primary human macrophages (Cervero et al., 2012). These class I myosins are monomeric motors that can bind actin filaments via a motor domain and plasma membrane via positively charged tail domains. Class I myosins include long-tailed (Myo1e and Myo1f, containing proline-rich and SH3 domains in their tails) and short-tailed myosins, whose tails consist only of the membrane-binding regions (McConnell and Tyska, 2010). In mammalian and yeast cells, class I myosins preferentially associate with Arp2/3-nucleated branched actin networks and nearby membranes in structures such as endocytic actin patches and phagocytic cups (Barger et al., 2019; Sirotkin et al., 2005). While Myo1e was previously detected in invadopodia of v-src-transformed fibroblasts and podosomes of RAW macrophages (Ouderkirk and Krendel, 2014; Zhang et al., 2019), the specific functions of Myo1e and Myo1f at podosomes and their contributions to macrophage migration have not been elucidated.

In the present study, we have interrogated the potential functional roles of Myo1e/f in podosome dynamics and macrophage migration using human macrophages differentiated from primary blood monocytes as well as murine bone-marrow derived macrophages (BMDMs) from the wild-type and Myo1e/Myo1f single and double knockout (dKO) mice (Barger et al., 2019; Kim et al., 2006; Krendel et al., 2009). Human cells readily form podosomes in large numbers, which facilitates statistical analysis, while murine knockout macrophages allow podosome analysis in the complete absence of our proteins of interest. Therefore, combining these two approaches enabled a comprehensive analysis of myosin I functions. Using both Myo1e/f depletion by siRNA and Myo1e/f knockout, we show that these motors specifically regulate podosome size, turnover, and lateral mobility. These abnormal podosome dynamics are associated with reduced macrophage migration, highlighting the role for Myo1e/f in macrophage motility.

Moreover, we find that Myo1e and Myo1f, but not other class I myosins tested, localize to the ventral surface of the podosome core, a region that in macrophages also contains the transmembrane proteins matrix metalloproteinase MT1-MMP (El Azzouzi et al., 2016) and the GPI-linked membrane-bound DNase X (Pal et al., 2021), as well as the hyaluronan receptor CD44 in osteoclasts (Chabadel et al., 2007). In line with its restricted localization underneath the podosome core and its specific components, we propose to name this part of the podosome architecture as the podosome “base”.

## Results

### Myo1e and Myo1f are enriched at the podosome base in macrophages derived from primary human monocytes

First, we examined the endogenous Myo1e and Myo1f localization at podosomes of human macrophages by immunofluorescence using myosin isoform-specific antibodies (**Fig. 1**). Both myosins localized to the F-actin rich podosome core labeled with phalloidin (**Fig. 1A-F, J-O**), but appeared concentrated at the lower part of the podosome core, close to the ventral surface. We hereby refer to this region as the podosome “base” (**Fig. 1G-I, P-R**, green). Analysis of fluorescence intensities using the Poji macro (Herzog et al., 2020) confirmed that the peaks of F-actin and Myo1e/f intensity all align along the central axis of podosomes and are equally wide, however, the signals for Myo1e/f show a maximum intensity in the confocal Z-layer between 0.5 – 1.0 µm above the substrate, while F-actin peaks between 1.0 – 1.5 µm (**Fig. 1S-V**). Like the endogenous proteins, exogenously expressed EGFP-tagged Myo1e and Myo1f also localized to podosomes (**Fig. 2C-E, Ć-É).** In contrast, none of the fluorescently tagged (EGFP or mEmerald) short-tailed class I myosins (Myo1a,b,c,d,g) were enriched at podosomes (**Supp.** Fig. 1).

**Figure 1.**
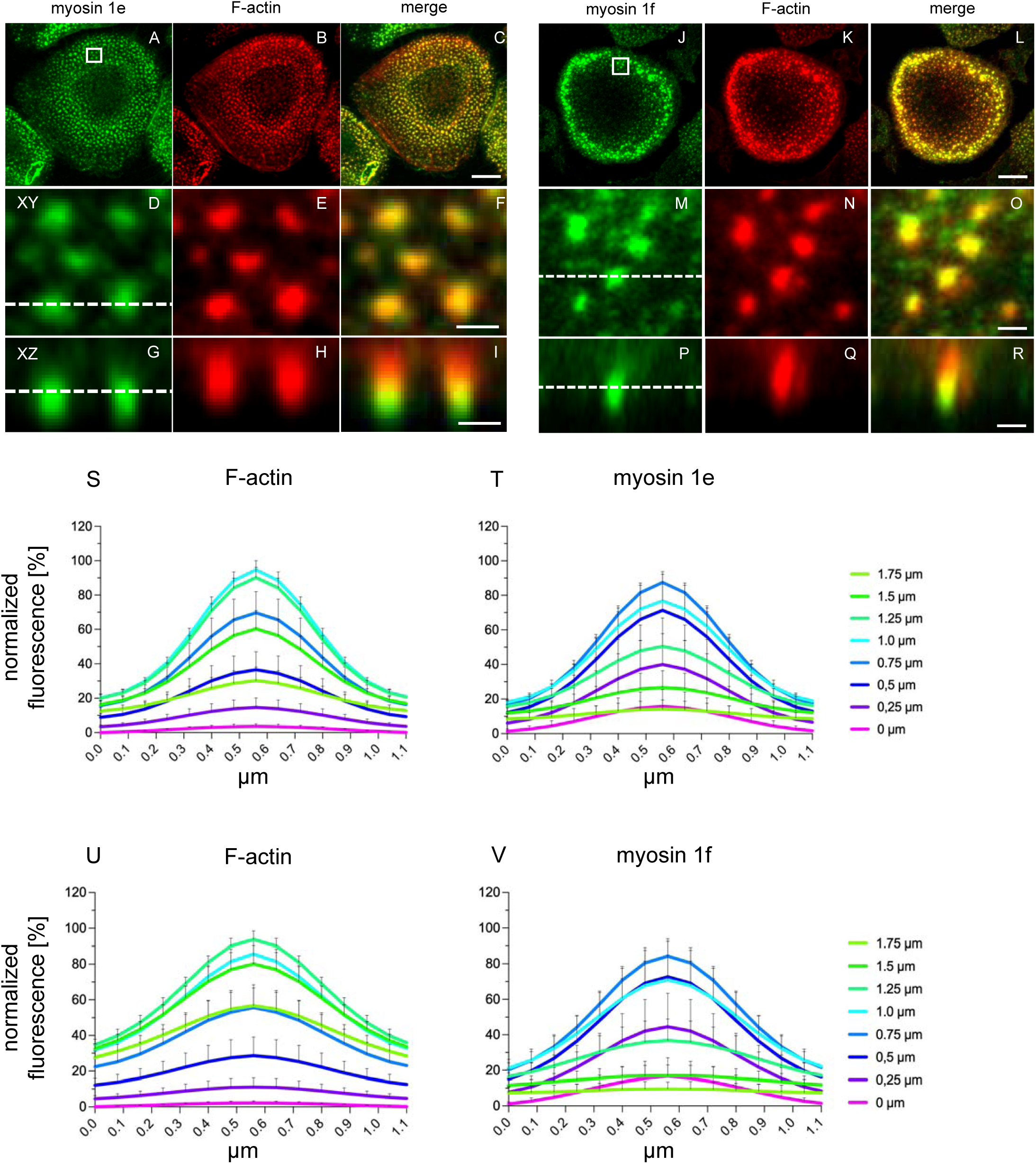
Endogenous Myo1e and Myo1f localize to the base of macrophage podosomes. (A-R) Confocal micrographs of primary human macrophages stained for endogenous Myo1E (A,D,G) or Myo1f (J,M,P), and for F-actin using Alexa-Fluor-568 phalloidin, to label podosome cores (B,E,H,K,N,Q), with respective merges (C,F,I,L,O,R). White boxes in (A,J) indicate regions of interest shown in (D-I, M-R), with respective XY (D-F, M-O) or XZ views (G-I, P-R). Note localization of Myo1e and Myo1f to the podosome base, a region mostly beneath the F-actin-rich podosome core. Scale bars: 10 µm for (A-C, J-L), 1 µm for (D-I, M-R). (S-V) Analysis of Myo1e and Myo1f localization at podosomes. Radial fluorescence intensity profiles of podosomes stained for F-actin (S,U) and Myo1e (T) or Myo1f (V). Profiles were calculated using PoJi, a Fiji-based tool (Herzog et al., 2020), and represent the mean ± s.e.m. of all podosome profiles imaged at indicated confocal z planes, with 0 µm set as the most ventral F-actin signal of podosomes; n=1411 podosomes, per z section, collected from three different cells (S, T) and n=1312 podosomes, per z section, collected from three different cells, (U, V). Note that Myo1e and Myo1f at podosomes show the maximum fluorescence intensity at z-plane of 0.75 µm, whereas F-actin peaks at z-planes of 1.0 – 1.25 µm.

**Figure 2.**
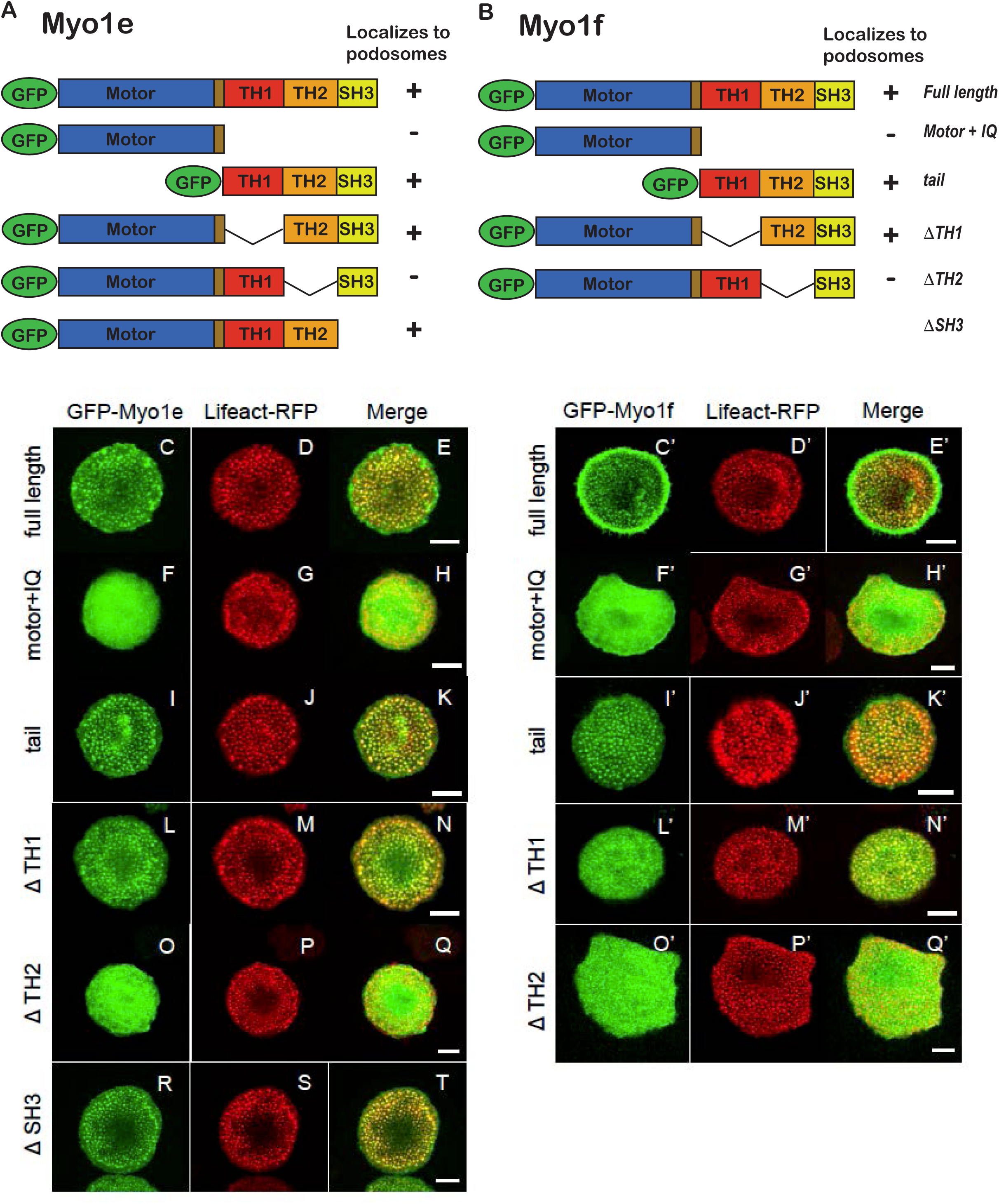
Localization of Myo1e and Myo1f to podosomes requires their TH2 domains. (A,B) Domain structure of Myo1e (A) and Myo1f (B). Both myosins contain an N-terminal motor domain, followed by a neck domain (containing an IQ motif) that binds calmodulin or other light chains, and a C-terminal tail, containing lipid-binding tail homology-1 (TH1), proline-rich tail homology-2 (TH2), and Src homology-3 (SH3) domains. Presence/absence of the respective EGFP-tagged expression constructs at macrophage podosomes is indicated by “+/-”. (C-T, C’-Q’) Confocal micrographs of primary human macrophages overexpressing GFP-fused full length or partial constructs of Myo1e (C,F,I,L,O,R) or Myo1f (C’,F’,I’,L’,O’), as indicated, with co-expression of lifeact-RFP, to label podosome cores (D’,G’,J’,M’,P’,S’,D’,G’,J’,M’,P’), with respective merges. Note absence from podosomes of constructs lacking the TH2 domain (“motor+IQ”, Δ”TH2”). Scale bars: 10 µm.

These data show that both endogenous and overexpressed forms of Myo1e and Myo1f localize to macrophage podosomes. Indeed, both motors are specifically enriched at the ventral side of podosome cores, which we name the podosome base. This localization is close to the ventral plasma membrane, which is likely correlated with the ability of both myosins to bind membrane lipids (Feeser et al., 2010; McIntosh and Ostap, 2016).

### The TH2 domains determine localization of Myo1e and Myo1f to podosomes

Myo1e and Myo1f contain a motor/head domain that binds ATP and F-actin, a neck domain (IQ) that binds calmodulin, and a tail, containing a lipid-binding tail homology-1 (TH1), proline-rich tail homology-2 (TH2), and Src homology-3 (SH3) domains (**Fig. 2A,B**). We have previously demonstrated that the TH2 domain of Myo1e was necessary for its localization to podosomes, and that the TH1 domain promoted its membrane enrichment/ventral localization (Ouderkirk and Krendel, 2014; Zhang et al., 2019). In contrast, localization requirements have not been previously examined for Myo1f. By overexpressing EGFP-tagged domain deletion mutants of Myo1e and Myo1f together with the RFP-labeled F-actin marker Lifeact, we tested the requirements for specific protein domains in localization of both myosins using live cell imaging to avoid fixation artifacts. We found, for both Myo1e and Myo1f, that constructs containing only the N-terminal motor and IQ domains (“motor + IQ”) (**Fig. 2 F-H, F’-H’**) were not enriched in podosomes, while constructs containing only the C-terminal half comprising the TH1, TH2 and SH3 domains (“tail”) were enriched in podosome cores (**Fig. 2 I-K, I’-K’**). Further dissection of the C-terminal tails showed that constructs lacking the TH2 domain (**Fig. 2O-Q, O’-Q’**) failed to localize to podosomes. In contrast, deletion of the TH1 domain (Fig. 2L-N, L’-N’) or of the SH3 domain (**Fig. 2 R-T**) did not interfere with their localization at podosomes. Collectively, these results show that the TH2 domains of Myo1e and Myo1f are required for the localization of both myosins to macrophage podosomes.

### Myo1e and Myo1f regulate size, lifetime and lateral mobility of podosomes

To test how Myo1e/f contribute to the regulation of podosome organization and dynamics, we examined the effects of the knockdown of these proteins in human macrophages, using two distinct siRNAs for each myosin to control for potential off-target effects (**Suppl. Fig.2A,B**). Analysis of live cell imaging videos of human macrophages expressing Lifeact-RFP to label podosome cores showed that Myo1e knockdown increased the size of podosome cores by ∼ 50% (0.38 ± 0.07 µm^2^ for controls, and values ranging from 0.56 ± 0.17 µm^2^ to 0.57 ± 0.2 µm^2^ for respective depletions) **(Fig. 3A)**, while treatment with all siRNAs led to reduced podosome lifetime by ∼30% (7.2 ± 3.1 min for controls, and values ranging from 4.9 ± 1.5 min to 5.4 ± 1.5 min for respective depletions) (**Fig. 3B**). Further analyses showed that the lateral displacement of podosomes during their lifetime was also reduced in these cells by ∼20% (1.0 ± 0.29 µm for controls, and values ranging from 0.77 ± 0.17 µm to 0.87 ± 0.23 µm for respective depletions) (**Figure 3C).**

**Figure 3.**
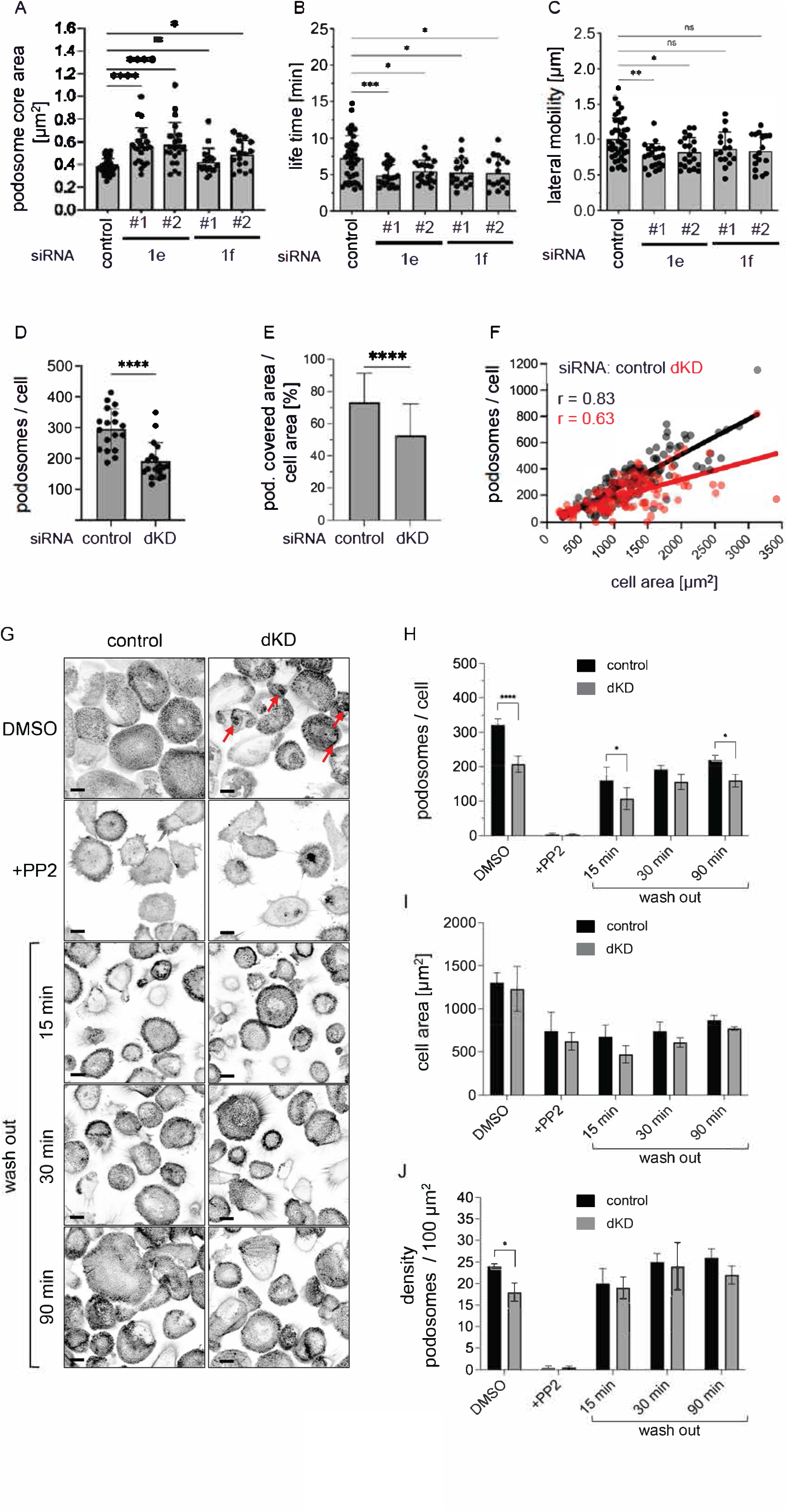
Myo1e/f regulate podosome size, lifetime, and lateral movement, but not reformation. (A-C) Analysis of podosome core area (A), life time (B) or lateral mobility (C) in macrophages treated with each time two individual siRNAs for Myo1e or Myo1f Ordinary one-way ANOVA, ****P < 0.0001, ***P < 0.001, **P < 0.01, *P < 0.05, ns = statistically not significant; at least 17 cells / condition were evaluated, collected from 3different donors. (D-F) Analysis of podosome numbers per cell, podosome-covered cell area. (D-F) Analysis of the number of podosomes per cell (D), the percentage of the area covered by podosomes relative to the area of the whole cell (E) and correlation analysis between the amount of podosomes formed by a specific cell and the respective cell area (F) in macrophages depleted simultaneously (dKD) for both Myo1e and Myo1f. Unpaired t-test, ****P < 0.0001; 18 cells / condition, collected from 3 different donors (D). Unpaired t-test, **P < 0.01; at least 120 cells / condition were evaluated, collected from three different donors (E). 120 cells / condition, collected from 3 different donors (F). (G-J) Podosome reformation assay to analyze the ability of podosomes to reform over time (15, 30, 90 min) after PP2-mediated dissolution, in macrophages simultaneously depleted for both Myo1e and Myo1f (dKD), with DMSO treated cells as controls. F-actin of podosome cores was stained with Alexa488-coupled phalloidin, and podosomes are visible as µm-sized black dots within cells. Red arrows in DMSO controls indicate large F-actin aggregates forming after dKD (G); Scale bars = 10 µm. ImageJ macro-based evaluation of the number of podosomes per cells (H), cell area (I) and related podosome density (J). Multiple unpaired t-test; ****P < 0.0001, *P < 0.05. At least 120 cells / condition were evaluated, collected from 3 different donors.

To further probe the potential influence of Myo1e and Myo1f on podosome genesis, we performed podosome reformation assays, which involve the complete disassembly of podosomes by the Src tyrosine kinase inhibitor PP2, followed by subsequent washout and podosome reassembly (Cervero et al., 2013). Because single knockdowns did not show clear effects on podosome reformation (not shown), we tested double knockdowns of Myo1e and Myo1f together. At time point 0 (no addition of PP2), human macrophages depleted for both Myo1e and Myo1f (**Suppl. Fig. 2B**) showed a reduction of podosome numbers by 35% (296 ± 66 podosomes/cell for controls, and 191 ± 61 podosomes/cell for double depletions) **(Fig. 3D)**, which was accompanied by a similarly reduced podosome covered area, compared to controls **(Fig. 3E).** Plotting cells by the number of podosomes in relation to cell area further showed that depletion of Myo1e/f led to a reduced correlation of cell size and podosome number, indicating that a smaller fraction of the cell area was covered by podosomes (r=0.83 for control cells vs. r=0.63 for Myo1e/f depleted cells) **(Fig. 3F)**. Similarly to individual knockdowns (**Fig. 3A**), double knockdown cells also showed an increase in podosome core size (**Suppl. Fig. 2C**). In addition, we observed increased formation of larger actin structures, possibly corresponding to podosome clusters or enlarged podosomes, in double depleted cells **(Fig. 3G).** At different time points after PP2 washout (15 min, 30 min, 90 min), Myo1e/f depleted cells showed reduced numbers of podosomes, compared to controls (**Fig. 3H**). However, as double-depleted cells also showed reduced cell size during the wash out (**Fig. 3I**), the density of podosomes (number of podosomes / 100 µm^2^) did not change significantly during the reformation (**Fig. 3J**).

Collectively, these data show that while Myo1e and Myo1f do not appear to play a direct role in podosome assembly, they affect the size, lifetime, and lateral mobility of podosomes. They also influence the number of podosomes per cell.

### Myo1e and Myo1f regulate podosomes and 2D/3D cell migration in murine macrophages

Having established the influence of Myo1e and Myo1f knockdown on podosomes in human macrophages, we next wanted to verify and expand on key findings using murine knockout macrophages. Similar to the results in knockdown human macrophages, double knockout of Myo1e/f led to increased size of podosomes **(Fig. 4A,B)**, which was associated with an increase in podosomal F-actin, as determined by fluorescence intensity measurements **(Fig. 4С**). Moreover, cells with single knockout of Myo1f showed a similar trend towards larger podosomes and increased F-actin content (**Fig. 4B,C**). While podosome cores did appear more clumped, the circularity of these structures was not significantly changed among the cells tested **(Fig. 4D**).

**Figure 4.**
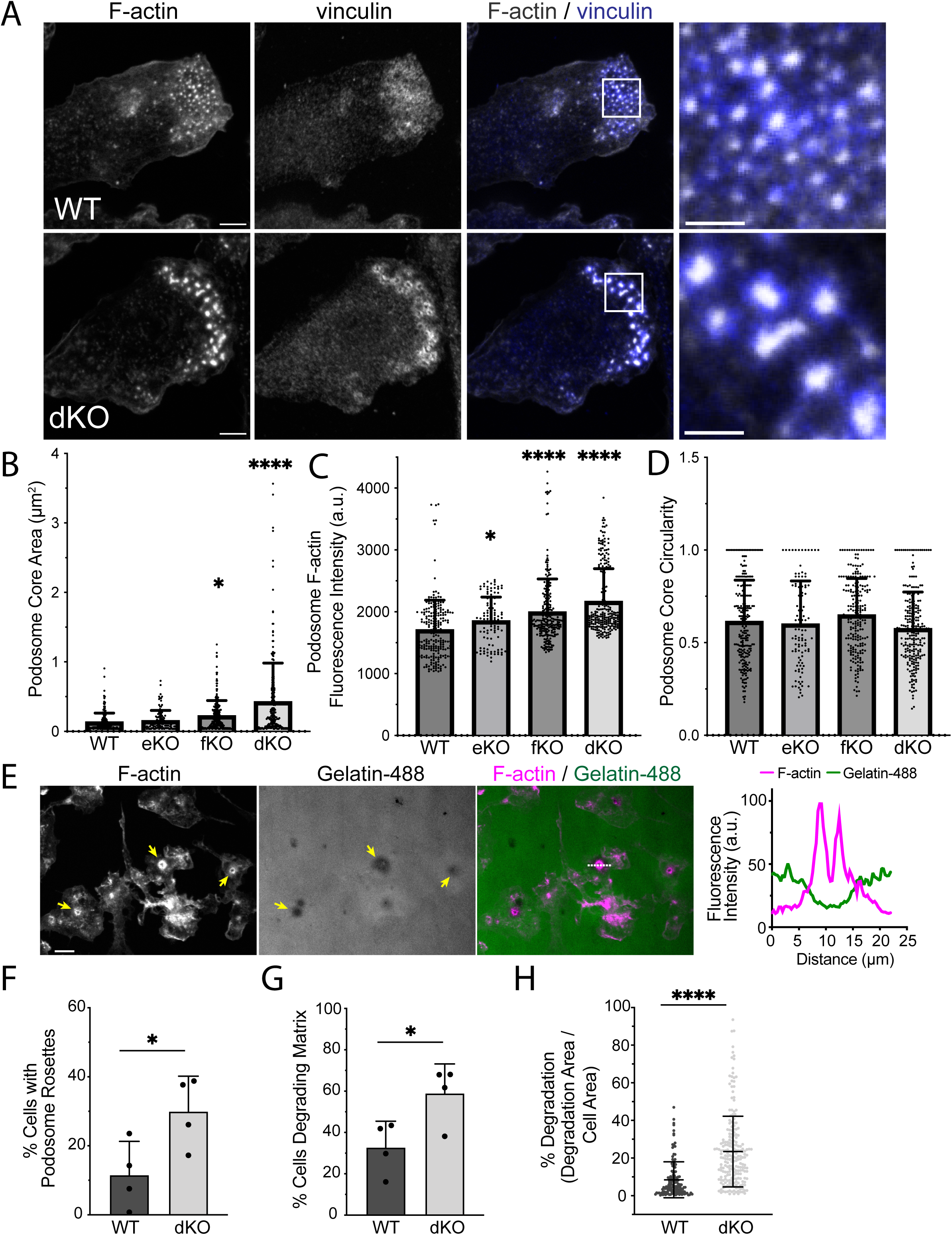
KO of Myo1e and Myo1f results in formation of podosome aggregates and increases matrix degradation. A, BMDMs derived from Myo1e/f dKO mice contain podosome clumps and aggregates. Macrophages were fixed and stained with phalloidin and anti-vinculin. (B,C,D) Graphs comparing (B) area, (C) fluorescence intensity, and (D) circularity of F-actin podosome cores of WT, eKO, fKO, and dKO BMDM podosomes (n = 100-250 podosomes measured in 10-20 cells). Data represents one representative experiment conducted in triplicate; each graph displays mean+SD and individual data points. * p<0.05, **** p<0.0001, compared to the WT by one-way ANOVA. B, The average area of podosome cores is increased in the Myo1f KO and Myo1e/f dKO BMDMs. C, Podosome mean F-actin intensity is higher in BMDM lacking Myo1e, Myo1f, or both Myo1e and Myo1f than in the WT cells. D, Podosome circularity is similar in all BMDM types. (E-H), gelatin matrix degradation by BMDMs. E, areas of matrix degradation are associated with F-actin-containing podosomes. F-G, Fraction of BMDMs containing podosome rosettes (F) or associated with the areas of matrix degradation (G) is higher in the dKO BMDM cultures. (N = 138 WT cells and 219 dKO cells pooled from 4 independent experiments). H, the area of matrix degradation associated with individual cells is higher in the dKO cells. * P<0.05 compared to the WT; **** P<0.0001 compared to the WT.

To test the degradative ability of the murine podosomes, we utilized the fluorescent gelatin degradation assay. Murine-derived macrophages were plated on fluorescent-gelatin coated coverslips and activated to form podosomes. Podosome-mediated degradation was confirmed by counterstaining with phalloidin (**Fig. 4E**). Compared to control macrophages, dKO cells exhibited podosome rosettes more frequently with a higher incidence of matrix degradation (**Fig. 4F,G**). Quantification of degraded area per cell area further showed that dKO macrophages exhibited increased matrix degradation **(Fig. 4H**).

Since podosome-mediated matrix degradation is required for macrophage invasion in a 3D context (Van Goethem et al., 2010), we tested the ability of Myo 1e/f dKO cells to migrate in 3D Matrigel. Surprisingly, in spite of the increased matrix degradation in 2D (**Fig. 4**), the number of dKO cells that migrated through Matrigel in 3D (**Fig. 5A**), as well as the maximum migration depth (**Fig. 5B**) were reduced compared to the WT cells. This finding may reflect the fact that macrophage 3D invasion is a complex process that relies not only on matrix degradation, but also on cell motility and adhesion capacity, which may also be altered by the Myo1e/f dKO.

**Figure 5.**
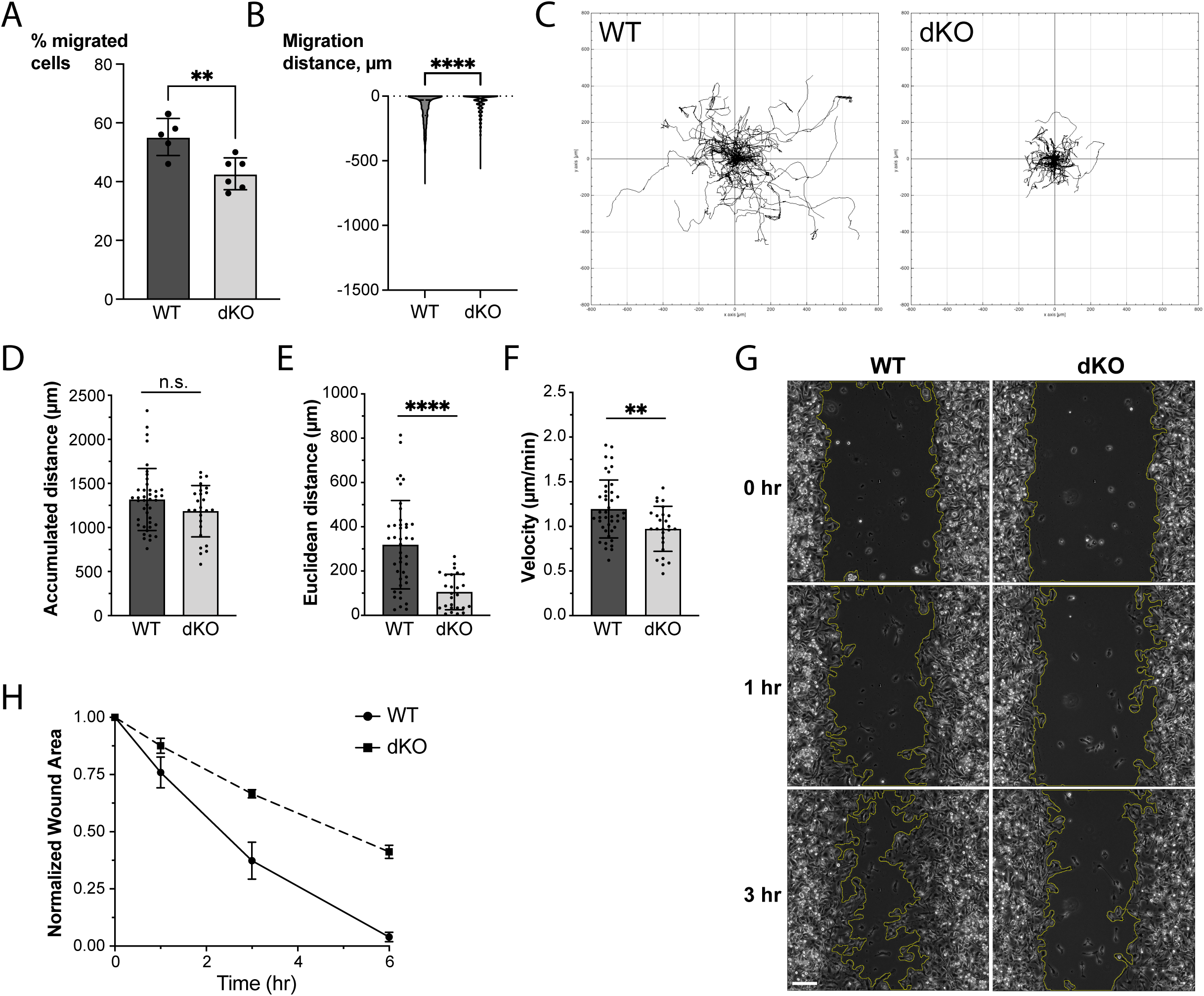
dKO BMDM interferes with BMDM migration. (A, B) The ability of dKO BMDMs to migrate into 3D Matrigel inserts was reduced compared to the WT cells. (A) The percentage of migration (dot) of BMDMs from 5 WT and 6 dKO mice is represented with the mean (bar) and SD. (B) The distribution of the distances migrated in Matrigel by individual cells obtained from 5 WT and 6 dKO mice is shown. (C) Tracks of WT and dKO macrophages randomly migrating on fibronectin-coated glass substrate. Quantification of tracks in (C) reporting (D) accumulated distance, (E) Euclidian distance, and (F) velocity. Representative results of 4 independent experiments, n = 42 WT and 26 dKO cells. (G). Representative images (phase contrast) of WT and dKO macrophages migrating into a scratch wound. Scale bar = 100 µm. (H) Quantification of wound area closure over 6 hrs (N=1 experiment, with data points representing 4 different FOVs). * P<0.05 compared to the WT; ** P<0.01 compared to the WT;**** P<0.0001 compared to the WT.

We thus additionally tested the effects of Myo1e/f on macrophage migration in 2D. To assess random migration in 2D, we tracked cell trajectories over the course of 20 hours (**Fig. 5C**). dKO macrophages were significantly less motile and appeared to have difficulty detaching from the substrate or sufficiently polarizing to form a leading edge (**Supp. Video 1**). Tracking the back-and-forth movement of the cell centroid of dKO macrophages showed that the total accumulated distance was lower in the dKO cells, although the difference was not significantly significant (**Fig. 5D**), however, little displacement was achieved from the starting point to the end point of each track (Euclidean distance, **Fig. 5E**). Furthermore, dKO cells moved at a significantly slower rate (**Figure 5F**). Finally, a wound-healing assay was used to measure the rate of directional cell migration into the wound (a scratch in the densely plated layer of cells). The rate of wound closure was significantly lower in the dKO cultures compared to WT cultures (**Figure 5 G-H**).

## Discussion

Class I myosins, which contain both actin-binding motor domains and membrane-binding tail domains, localize to actin-membrane contact sites and have been linked to vesicular trafficking, membrane deformation during clathrin-dependent endocytosis, cell migration, and regulation of cell adhesion (McIntosh and Ostap, 2016). Positively charged tail homology domains of these myosins (TH1 domains) promote association with phospholipids, in some cases exhibiting very specific lipid-binding preferences (Myo1c) (Hokanson and Ostap, 2006) and in others having broader phospholipid binding specificity (Myo1e) (Feeser et al., 2010). A subclass of class I myosins is formed by the so-called long-tailed class I myosins, whose tails include additional domains, such as the positively charged and proline-rich TH2 domain and the SH3 domain that can bind proline-rich domains (PRDs). These myosins are found in a variety of organisms, including yeast, amoeboid Protozoans, and vertebrates, with mammalian genomes encoding two long-tailed myosins, Myo1e and Myo1f (Kim and Flavell, 2008).

Cellular structures enriched in long-tailed class I myosins, including endocytic actin patches (Krendel et al., 2007; Manenschijn et al., 2019; Sirotkin et al., 2010), actin waves (Brzeska et al., 2014), and phagocytic actin cups (Barger et al., 2019), have several characteristic features. These structures are assembled in the regions of the plasma membrane enriched in PI(3,4)P_2_, PI(4,5)P_2_ or PI(3,4,5)P_3_, consist of branched, Arp2/3-nucleated actin networks, and contain multiple SH3- and PRD-containing proteins, including members of the WASp family of Arp2/3 activators. Podosomes and invadopodia share many of the same characteristics – their assembly is triggered by the clustering of specific phospholipids and involves recruitment of multiple SH3- and PRD-containing proteins and assembly of branched actin via WASp/N-WASp-activated Arp2/3-mediated nucleation (Linder et al., 2023).

We have previously shown that Myo1e localizes to the ventral podosome surface in src-transformed fibroblasts, RAW mouse macrophages, and A375 melanoma cells (Ouderkirk and Krendel, 2014; Zhang et al., 2019) and that both Myo1e and Myo1f can be identified in podosome-enriched macrophage fractions by proteomic analysis (Cervero et al., 2012). We have also previously observed both Myo1e and Myo1f in phagosome-associated podosomes (Barger et al., 2019).

In this paper, we have dissected Myo1e and Myo1f localization to macrophage podosomes in more detail and identified defects in podosome dynamics and macrophage migration that are associated with their depletion. We find that both Myo1e and Myo1f, but not other class I myosins, localize to the region of podosomes that is close to the ventral plasma membrane of cells. Previously identified podosome components that share a similar localization are the matrix metalloproteinase MT1-MMP (El Azzouzi et al., 2016), DNase X (Pal et al., 2021) and the hyaluronan receptor CD44 (Chabadel et al., 2007). All of these proteins are attached to at least one leaflet of the ventral plasma membrane, either through a transmembrane domain or through a GPI anchor, in case of DNaseX, whereas Myo1e/Myo1f interact with the membrane phospholipids via their tail domains. In line with this restricted localization beneath the podosome core and its specific components, we propose to label this part of the podosome architecture as the podosome “base”. The podosome base would thus join the current trifecta of podosome core, ring and cap as an additional substructure with a defined molecular composition and function.

Similarly to the previous localization studies (Barger et al., 2019; Ouderkirk and Krendel, 2014; Zhang et al., 2019), Myo1e/f localization to macrophage podosomes depended on the TH2 domain. None of the short-tailed class I myosins we tested, which lack TH2 and SH3 domains, localized to podosomes, further highlighting the importance of the myosin C-terminal tail region for podosome localization. Both TH1 and TH2 domains of Myo1e and Myo1f are highly positively charged, which presumably promotes their interactions with membrane phospholipids, however, the TH1 domain does not appear to be required for recruitment to podosomes. In addition to carrying positive charges, the Myo1e/f TH2 domains may be able to interact with SH3 domain-containing proteins via the proline-rich motifs in the TH2 domain and may also bind actin filaments in an ATP-independent fashion similarly to the long-tailed amoeboid myosins I (Rosenfeld and Rener, 1994), although the actin-binding ability has not been confirmed for mammalian Myo1e/f TH2 domains. Whether these activities further contribute to their role in podosome localization or dynamics remains to be determined.

Simultaneous knockout of Myo1e/f in mouse macrophages or knockdown of these myosins in human macrophages resulted in formation of enlarged podosomes or podosome aggregates concomitant with a reduction in the number of podosomes per cell. Unlike the overexpression of the TH1-TH2 domains of Myo1e, which blocks macrophage podosome formation and suppresses gelatin degradation by cancer cells (Zhang et al., 2019), Myo1e/f depletion did not result in the loss of podosomes or their substrate-degrading activity. It is possible that the overexpression of the podosome-localized TH1-TH2 fragment may act in a dominant-negative fashion by disrupting additional protein-protein and protein-lipid interactions within nascent podosomes and further impairing other cellular functions. Myo1e/f depletion does not have this effect on the overall podosome assembly but instead disrupts specific myosin-dependent steps in podosome dynamics and reorganization.

We also find that individual knockdowns of Myo1e or Myo1f in human macrophages lead to reduction in podosome lifetime and lateral mobility. Podosome reformation experiments showed that the decreased lifetime is not based on the reduced de novo formation of podosomes but likely on higher turnover rates. Myo1e/f activity may contribute to podosome dynamics and organization via several possible mechanisms. We have previously shown that macrophages lacking Myo1e/f exhibit reduced membrane tension (Barger et al., 2019). Artificially increasing membrane tension by osmotic swelling causes macrophage podosomes to disassemble, while artificially decreasing membrane tension causes podosomes to cluster (Rafiq et al., 2019). This could explain our observations in Myo1e/f dKO or dKD cells, where podosomes are enlarged or form clusters. These findings are also reminiscent of our observations on the effects of Myo1e/f KO on phagocytic actin waves/phagocytic podosomes, where the lack of Myo1e/f was associated with excessive branched actin assembly and the replacement of individual phagocytic podosomes with large actin aggregates (Barger et al., 2019). In addition, by mediating actin-membrane interactions, Myo1e/f may also regulate the angle of actin filament-membrane contacts or modulate bending or compressive forces on actin filaments (Jasnin et al., 2022), which in turn would affect the density of branched actin networks (Li et al., 2022; Risca et al., 2012) or even promote actin debranching (Pernier et al., 2020). Indeed, a recent study has identified a role for the motor activity of a class I myosin in promoting formation of sparser/less densely branched actin networks around beads coated with Arp2/3-activating proteins (Xu et al., 2024). Actin-based motility of Myo1e/f, possibly coupled with PIP-dependent anchoring to the plasma membrane, could also contribute to the observed roles of Myo1e/f in lateral mobility of podosomes by moving actin networks relative to the membrane or moving membrane PIPs relative to the underlying actin structures.

We have also found that Myo1e/f dKO macrophages exhibit slower cell migration in both 2D and 3D settings, which is in line with some of the previous observations regarding the effects of Myo1e/f depletion on cell motility (Garone et al., 2023; Giron-Perez et al., 2020; Kim et al., 2006; Martinez-Vargas et al., 2023; Navines-Ferrer et al., 2019; Tanimura et al., 2016). The fact that Myo1e/f dKO macrophages are more degradative could also stem from their reduced motility, i.e. because KO macrophages are moving less, they are degrading more of the area beneath them. Given that recruitment of podosomes to the leading edge plays an important role in macrophage symmetry breaking and migration initiation (Cervero et al., 2018), the disruption of podosome organization and dynamics could explain reduced migration in the absence of Myo1e/f. Whether the observed defects in podosome dynamics represent the sole reason for reduced cell migration upon loss of Myo1e/f requires further study since Myo1e/f may also promote lamellipodial protrusion (Tanimura et al., 2016) or modulate intracellular signaling (Giron-Perez et al., 2020; Heim et al., 2017). Overall, our findings suggest that Myo1e/f are crucial not only for podosome dynamics but also for coordinating the balance between adhesion, degradation, and motility required for efficient macrophage migration.

## Acknowledgements

We thank Frank Bentzien (UKE transfusion medicine) for buffy coats, Andrea Mordhorst and Sharon Chase for expert technical assistance, the UKE microscopy facility (umif) for help with microscopy and image analysis, and Martin Aepfelbacher for continuous support. Work on podosomes in our labs is supported by Deutsche Forschungsgemeinschaft (LI925/13-1 to SL), the Université Toulouse III -Paul Sabatier (UT3) (to RP) and by National Institute of General Medical Sciences (Award Number R01GM138652 to MK).

The authors declare no competing financial interests.

## Materials and Methods

### Expression constructs

EGFP-tagged and mEmerald-tagged Myo1e/f constructs as well as deletion constructs have been previously described (Barger et al., 2019; Bi et al., 2013). LifeAct-TagRFP was purchased from Ibidi (Martinsried, Germany).

### Antibodies and staining reagents

The following commercial antibodies were used: mouse monoclonal anti-vinculin (Sigma, H-VIN), mouse monoclonal anti-Myo1f (Santa Cruz, sc-376534), rabbit anti-Myo1e (RRID:AB_2909514) (Skowron et al., 1998), rabbit polyclonal anti-GAPDH (Proteintech, 10494-1-AP).

Secondary antibodies labeled with Alexa Fluor-488 and Alexa Fluor-568 and fluorescently-labeled phalloidin were purchased from Invitrogen. HRP-linked F(ab’)2 fragment donkey-anti rabbit (NA9340v) and HRP-linked sheep-anti mouse IgG (H + L) (NA931v) were purchased from GE Healthcare.

### Macrophage preparation

#### Mouse macrophages

All experiments involving mouse bone marrow isolation were performed according to the animal protocol approved by the Upstate Medical University IACUC. Myo1e and Myo1f KO mice were maintained on the C57Bl/6 background and all experiments used age- and sex-matched bone marrow preparations. BMDM isolation and differentiation has been previously described (Barger et al., 2022).

#### Human macrophages

Peripheral blood monocytes were isolated from buffy coats (obtained from University Medical Center Hamburg-Eppendorf, Hamburg, Germany) as described previously (Wiesner et al., 2010). The buffy coats were purified using Lymphocyte Separation Medium (LSM), a Ficoll-based medium. Then, the monocytes were positively selected for CD14. Cells were then differentiated into macrophages.

#### Cell Culture

BMDMs were maintained in DMEM supplemented with 10% FBS, 1% antibiotic/antimycotic (Life Technologies), and 20 ng/ml recombinant murine M-CSF (Biolegend; 576404). Human macrophages were cultured in RPMI medium supplemented with 20% autologous serum, penicillin, and streptomycin at 37 °C, 5% CO_2_. Transfections were performed using the Neon Electroporation System (Thermo Fischer Scientific).

#### 3D Cell Migration

Matrigel was slowly thawed on ice, deposited into the Transwell inserts (8µm-pore size polyester membrane, Corning 353097) in a 24-well plate (100 µl of 10mg/ml Matrigel per insert), and allowed to polymerize at 37°C. Matrices were rehydrated overnight with RPMI without FBS. Differentiated BMDM were serum-starved for 3 hrs and added to the top of the Matrigel plug in RPMI medium supplemented with 2% FBS and 20 ng/ml mouse M-CSF (4×10^4^ to 5×10^4^ cells per insert). The bottom well was filled with RPMI medium containing 10% FBS and 20 ng/ml mouse M-CSF, and cells were allowed to migrate for 2 days (approximately 48 hrs). Brightfield images of cells at the surface and within the Matrigel were collected as z-stacks with 30 µm intervals using a Nikon Eclipse TE2000-E2 multimode TIRF microscope equipped with PRIME-95B cMOS camera (Photometrics). Quantification was performed as in (Van Goethem et al., 2010). Two independent sets of experiments were performed, using BMDM from a total of 5 individual WT mice and 6 dKO mice. We used the mean percentage of migrated cells (relative to total cell number) from 2 to 3 inserts per mouse as technical replicates.

### Immunostaining

#### Mouse macrophages

Cells were fixed using fresh 4% paraformaldehyde/PBS for 15 min. After washing away fixative, cells were permeabilized in 0.1% Triton X-100/PBS for 3-5 min. Cells were blocked for 30 min at room temperature with 5% normal goat serum/3% BSA dissolved in PBS with 0.05% Tween-20 and 0.1M glycine. Cells were exposed to primary antibodies at the appropriate dilutions for 1 h at room temperature. Cells were then washed three times for 5 min with PBS. Secondary antibodies and fluorescent phalloidin were then applied for 30 min at room temperature. Cells were then washed again for three times, 5 min each before mounting with Prolong Diamond Antifade Mountant (Invitrogen). NucBlue Fixed Cell ReadyProbes Reagent (Invitrogen) was used according to the manufacturer’s instructions.

#### Human macrophages

2-weeks old macrophages were seeded on 12-mm glass coverslips (∼7 x 10^4^ cells per coverslip) and incubated overnight at 37 °C, 5% CO2, and 90% humidity. To fix, the coverslips were dipped for 1-3 seconds in glacial methanol (−20 °C) then placed in 3.7% formaldehyde for 10 min. The coverslips were then placed in PBS and stored at 4°C overnight. The cells were permeabilized for 5 min using 0.5% Triton X-100, followed by three washes with PBS containing 0.05 % of Triton X-100. Then, the samples were incubated for 1hr in blocking solution (2% BSA + 5% goat serum in PBS), followed again by three washes. They were then incubated with primary antibody solution for 2 hrs, followed by three washes. Finally, they were incubated for 1h with secondary antibody solution, which included a fluorescently-labelled phalloidin. After three PBS washes and a dip in dH_2_O, the coverslips were mounted on glass slides using Mowiol heated to 37°C. Primary antibodies were used at the following concentrations: anti-Myo1e (1:100), anti-Myo1f (1:100).

### Western blotting

Cells were lysed with RIPA buffer (150 mM NaCl, 1% Triton X-100, 0.5% sodium deoxycholate, 0.1% SDS, 50mM Tris–HCl (pH 8.0)) and vortexed. Equal amounts of protein samples were then mixed with 4× Laemmli sample loading buffer and examined by standard immunoblotting procedure using NuPAGE 4–12% Bis-Tris gels (Invitrogen, Thermo Fisher Scientific), iBlot2 dry blotting system (Thermo Fisher Scientific) and the above-mentioned primary and HRP-conjugated antibodies, as indicated. When needed, nitrocellulose membranes were mild-stripped by extensive washing with buffer (200 mM glycine, 3.5 mM SDS, 1% Tween-20, (pH 2.2)) before reblocking and reprobing membranes with primary and secondary HRP-conjugated antibodies. Protein bands were visualized by using Super Signal Pico or Femto kit (Pierce) and X Omat AR films (Kodak). Results were scanned and protein band intensities quantified with Fiji distribution of ImageJ (NIH, Bethesda, MD).

### Gelatin degradation assay

Gelatin was purchased from Sigma while Oregon-Green-488-labeled gelatin was purchased from Thermo Fisher, and gelatin-coated coverslips were prepared as previously described (Diaz, 2013). WT and dKO BMDM were plated on fluorescent gelatin and treated with 20 ng/mL IL-4 to induce podosome formation and ECM degradation (Cougoule et al., 2012). After 24 hrs, cells were fixed using fresh 4% paraformaldehyde/PBS for 15 min. After washing away fixative, cells were permeabilized in 0.1% Triton X-100/PBS for 3 min and stained with Alexa Fluor-568 phalloidin for 30 min. During washes, cells were counterstained with NucBlue and carefully mounted using Prolong Diamond Antifade Mountant. All plating and staining steps were performed under reduced lighting.

### Microscopy

#### Mouse macrophages

Live cell imaging to examine BMDM motility was performed on a True MultiColor Laser TIRF Leica AM TIRF MC system equipped with an Andor DU-885K-CSO-#VP camera and a 10X objective (Leica HCX PL FLUOTAR 10x/0.30NA PH1). Environmental conditions were maintained by an Okolab temperature- and CO_2_- control system. Spinning disk images were collected using a PerkinElmer UltraView VoX Spinning Disc Confocal system mounted on a Nikon Eclipse Ti-E microscope equipped with a Hamamatsu C9100-50 EMCCD camera, a 100X (1.4 N.A.) PlanApo objective, and controlled by the Volocity software.

#### Human macrophages

Imaging of Myo1e/f deletion constructs and Myosins 1 a/b/c/d/g was performed in a live cell set-up using a Visitron spinning disk microscope equipped as follows: Nikon Eclipse TiE microscope; 60X Apo TIRF Oil objective with NA 1.49; Photometrics Prime 95B camera; Yokogawa CSU W-1 spinning disk unit; VisiView software. Images of fixed samples stained for endogenous Myosins I e/f were acquired using a confocal laser scanning microscope (DMI 6000 with a TCS SP5 AOBS confocal point scanner; Leica) equipped with an oil-immersion HCX PL APO 63X NA1.4-0.6 objective. Where needed, Z-stacks were acquired at 0.25 μm intervals.

### Image analysis

#### Mouse macrophages

Fiji (Schindelin et al., 2012) was used to process and analyze all microscopy images. To make Z-projections, Z-stacks that displayed the podosomes were flattened using average intensity projections. The same stacks were used for all channels in an image. The contrast on the images were then enhanced to 0.1% and normalized. To generate intensity profiles, a podosome was selected, and line scans of fluorescent intensity were generated using a linear selection and the “plot profile” function in Fiji was used to obtain intensity values along the line. Wound healing analysis was performed using the MRI Wound Healing plugin, and tracking random motility was conducted using the Manual Tracking and Chemotaxis Tool plugins. Podosome core size measurements as well as areas of gelatin degradation measurements were accomplished using the Thresholding function in Fiji to generate ROI masks and measured using the Fiji Shape Descriptors function.

#### Human macrophages

Detailed analysis of several podosome profiles at different Z-positions was performed using the Fiji-based macro code termed ’Poji’ (Herzog et al., 2020). PP2 reformation assay and measurements of number of podosome per cell and podosome core areas were performed as previously described (Cervero et al., 2013). Tracking of podosome lifetime and podosome lateral mobility was performed using the Fiji-based Trackmate plugin (Ershov et al., 2022).

### Statistical analysis

Comparisons between WT and dKO BMDM were carried out using an unpaired two-tailed t-test for independent samples, with differences between genotypes considered statistically significant at p < 0.05. For multiple comparisons (WT, eKO, fKO, dKO), data were analyzed using a one-way ANOVA with Tukey’s post-hoc test, with statistical significance set at p-value < 0.05. For primary human macrophages, comparisons between controls and individual siRNAs were performed using ordinary one-way ANOVA with Dunnett’s multiple comparisons test, whereas comparisons between controls and Myo1e/1f double knockdown were performed using unpaired two-tailed t-test. For the podosome reformation assay, multiple unpaired t-tests were used. For quantification of knockdown efficiency by western blot, the intensities of specific proteins were first normalized for respective GAPDH intensities, then, after setting the control samples to 100%, the protein expression of Myo1e/1f (single and double knockdowns) was calculated relative to the control, and the resulting data analyzed by one sample t and Wilcoxon test. In all cases the statistical significance was set at p-value < 0.05. Statistical analyses and graphing were performed using GraphPad Prism software.

## Supplementary Figure legends

**Suppl. Figure 1.**
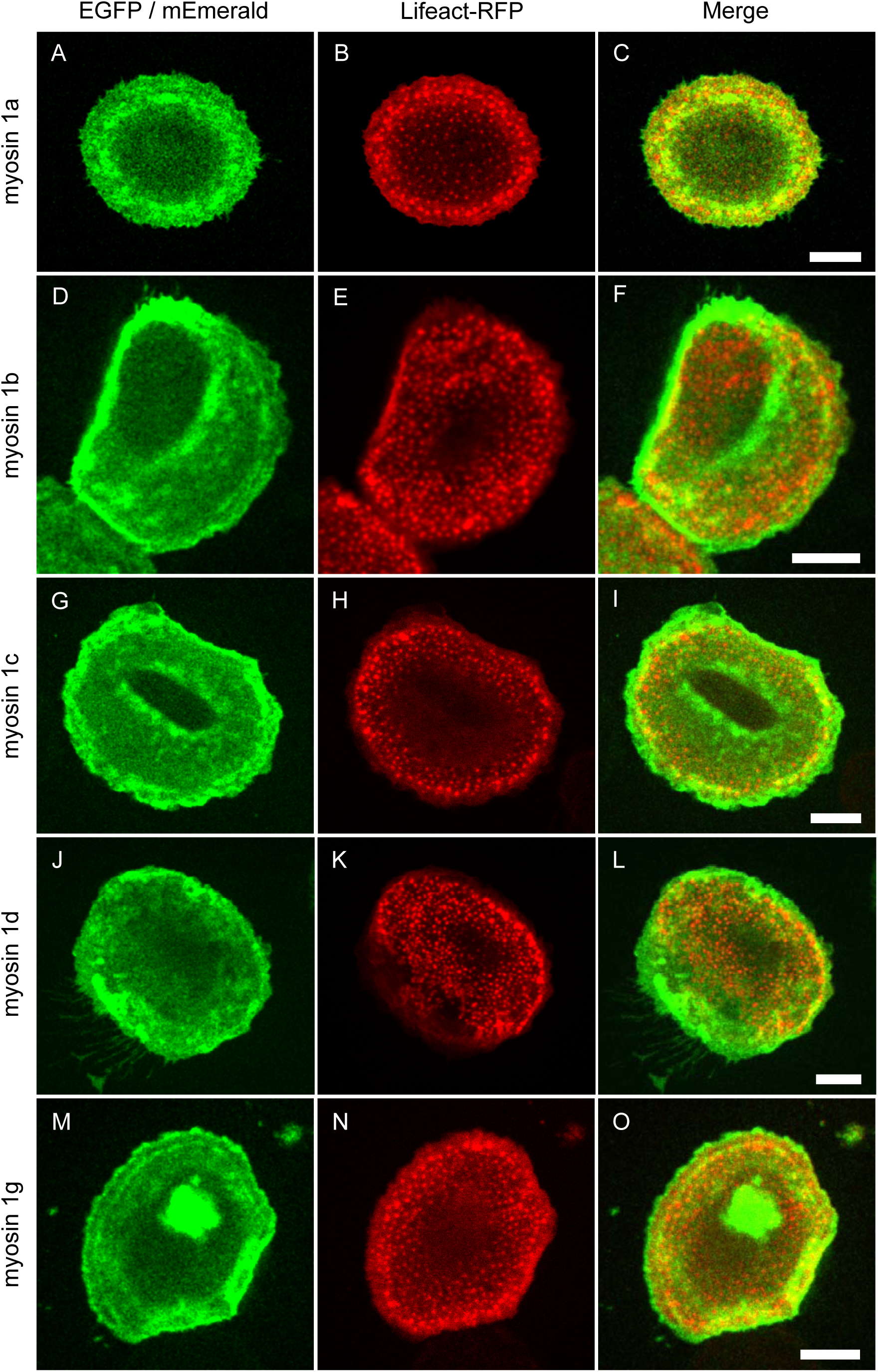
Short-tailed class I myosins do not localize to macrophage podosomes. Confocal micrographs of primary human macrophages overexpressing fluorescently labeled (EGFP or mEmerald) constructs of short-tailed class I myosins, including Myo1a (A), Myo1b (D), Myo1c (G), Myo1d (J), or Myo1g (M), and coexpressing lifeact-RFP, to label podosome cores (B,E,H,K,N), with respective merges (C,F,I,L,O). Scale bars: 10 µm.

**Suppl. Figure 2.**
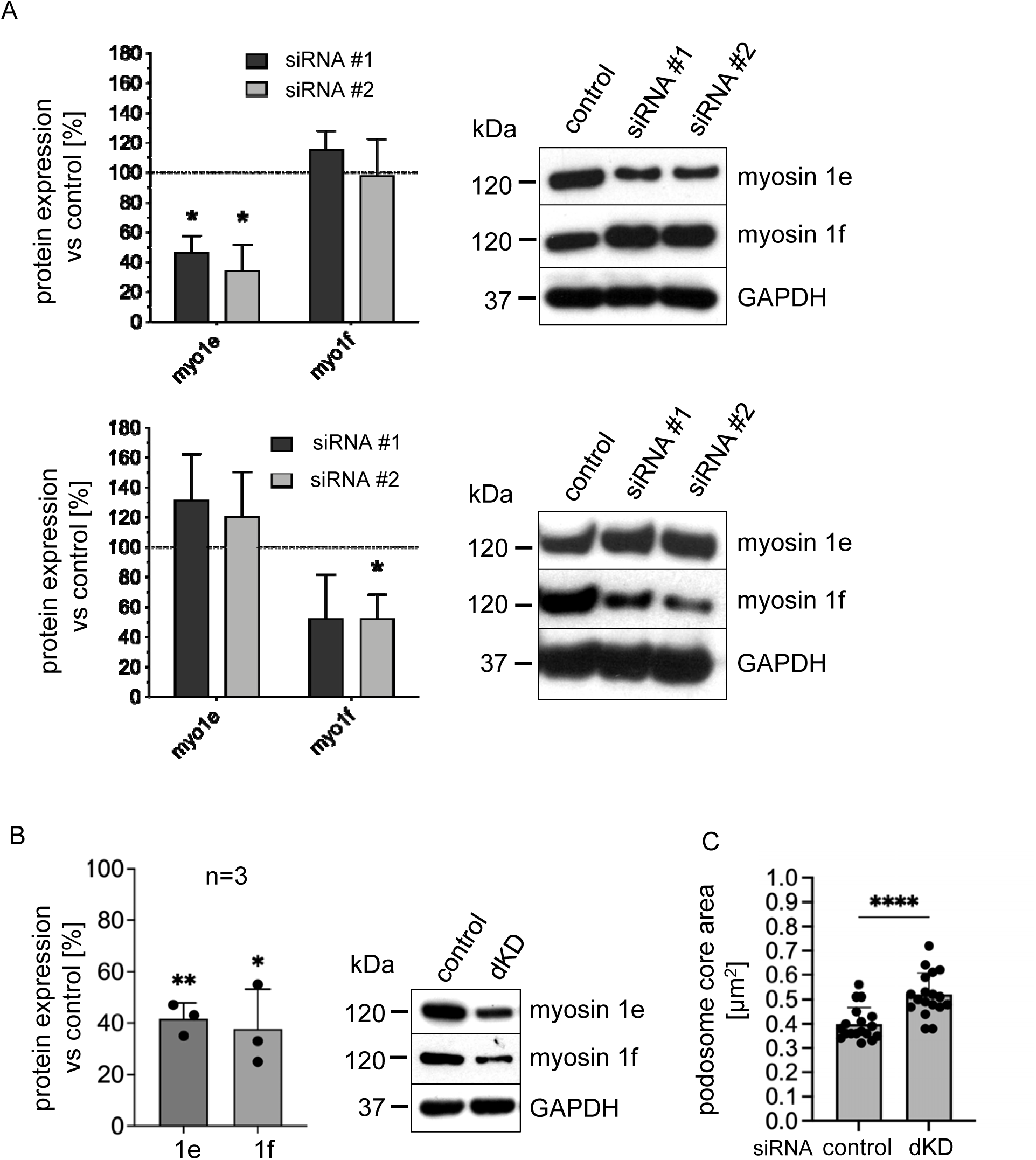
Quantification and effect of Myo1e/f knockdowns. (A) Knockdown of Myo1e and Myo1f with each time two independent siRNAs, with analysis of respective efficiency by Western blot. One sample t and Wilcoxon test, *P < 0.05; n=3 different donors. (B) Double knockdown (dKD) of both Myo1e and Myo1f, with analysis of respective efficiency by western blot. One sample t and Wilcoxon test, **P < 0.01, *P < 0.05; n=3 different donors. (C) Analysis of podosome core area by ImageJ macro in cells simultaneously depleted (dKD) for both Myo1e and Myo1f. Unpaired t-test, ****P < 0.0001; 18 cells / condition, from 3 different donors.

**Suppl. Video 1 (related to Figure 5). WT and dKO BMDM migrating on fibronectin-coated glass substrates.**

